# Fibroblast Growth Factor Signaling Instructs Ensheathing Glia Wrapping of *Drosophila* Olfactory Glomeruli

**DOI:** 10.1101/151662

**Authors:** Bing Wu, Jiefu Li, Ya-Hui Chou, David Luginbuhl, Liqun Luo

## Abstract

The formation of complex yet highly organized neural circuits requires interactions between neurons and glia. During the assembly of the *Drosophila* olfactory circuit, 50 olfactory receptor neuron (ORN) classes and 50 projection neuron (PN) classes form synaptic connections in 50 glomerular compartments in the antennal lobe, each of which represents a discrete olfactory information processing channel. Each compartment is separated from the adjacent compartments by membranous processes from ensheathing glia. Here we show that Thisbe, a fibroblast growth factor (FGF) released from olfactory neurons, particularly local interneurons, instructs ensheathing glia to wrap each glomerulus. The Heartless FGF receptor acts cell-autonomously in ensheathing glia to regulate process extension so as to insulate each neuropil compartment. Overexpressing Thisbe in ORNs or PNs causes over-wrapping of glomeruli to which their axons or dendrites target. Failure to establish the FGF-dependent glia structure disrupts precise ORN axon targeting and discrete glomerular formation.

**Significance Statement:** This research reports that reciprocal interactions between *Drosophila* olfactory neurons and ensheathing glia mediate the formation of neuronal compartments—groups of synapses that are packed into discrete structures called glomeruli that carry specific olfactory information. Ensheathing glia respond to a neuronal cue, the fibroblast growth factor (FGF) Thisbe, to pattern the boundaries of the nascent compartments. Neural compartments in turn require such glial barriers to separate themselves from neighboring compartments, so as to ensure the correct organization of the olfactory circuit. These findings highlight the importance of glia in the assembly and maintenance of neural circuits and the functions of FGF signaling in these processes.

## Introduction

Glia and neurons interact dynamically to coordinate the development and function of neural circuits. For example, *Drosophila* midline glia serve as guideposts that provide spatial cues to direct axonal pathfinding in the embryonic ventral nerve cord (1, 2). Neurons in turn provide signals that weave glia into the fabric of the nervous system. For instance, axonal Neuregulin-1 levels dictate the extent of Schwann cell myelination (3). A myriad of other biological processes in the developing and adult brain, including synapse formation (4-7), elimination (8, 9), and regulation of synaptic activity (10-13), all require extensive communication and cooperation between neurons and glia.

The *Drosophila* nervous system contains a diverse array of glial cell types (14), offering opportunities to discover new mechanisms for glia-neuron interactions. Ensheathing glia are a group of cells that insulate neighboring neuropil structures by extending their membranous processes along the outer surface of synaptic neuropil or their sub-compartments, without invading the inner part of the neuropil (15). It remains unknown how ensheathing glia establish such a barrier-like structure, what molecular signals orchestrate this process, and whether the glial barrier is essential for the integrity of the encircled neuropil.

The *Drosophila* antennal lobe provides an excellent experimental system for tackling these questions with high resolution. The antennal lobe is organized into 50 discrete neuropil compartments—the glomeruli wherein specific types of olfactory receptor neuron (ORN) axons and projection neuron (PN) dendrites form synapses—each of which is surrounded by ensheathing glia processes (15-17). In addition, the specific and stereotypic projection pattern of neurons in the antennal lobe (18, 19) renders the system convenient for exploring the potential neuronal disorganization caused by malformation of glial structures. Previous studies have utilized this system to find that in adults, ensheathing glia increase their process extension to injured axons (20), help clear degenerating axons (20), and strengthen excitatory interactions between surviving neurons (21). Here we study ensheathing glia in the antennal lobe formation and organization during development.

## Results

### Pupal Development of Antennal Lobe Ensheathing Glia

To study the development and function of antennal lobe ensheathing glia, we searched for genetic tools that can specifically label these cells during the pupal stage when the olfactory circuit in the antennal lobe is being assembled. We preselected a number of enhancer-GAL4 lines from the FlyLight GAL4 collection based on the published expression pattern in adult animals (22), and a few *MiMIC-GAL4* lines (23). We tested whether these GAL4 lines were active at different pupal stages. We found that at 96 hours after puparium formation (h APF), a late stage in antennal lobe morphogenesis, *SPARC-GAL4* (Fig 1D), *GMR56F03-GAL4* (Fig S1) (24), and *GMR10E12-GAL4* (Fig S1) flies expressed GAL4 predominantly in ensheathing glia. As indicated by the *UAS-mCD8GFP* reporter that labeled the cell membranes and processes in the presence of GAL4, these GAL4+ cells were located on the periphery of the antennal lobe and extended their processes to wrap around the antennal lobe as well as individual glomeruli within it, but with minimal invasion into each glomerulus. These are characteristic morphological features of ensheathing glia. We also used *UAS-nuclear-LacZ* to mark the nuclei of these GAL4+ cells (Fig 1E), and found that every LacZ+ nucleus around the antennal lobe (within 10-μm distance to the surface of the antennal lobe) was also positive for Repo, a glia marker in *Drosophila* (25), confirming that these GAL4+ cells are indeed glia. On average ~100 Repo+ cells were detected around the antennal lobe, and ~56 were LacZ+ (Fig 1F). Repo+/LacZ– cells are likely astrocytes and cortex glia present in the vicinity of ensheathing glia. To further distinguish these GAL4+ glia from astrocytes, we used the GAT antibody (26) to mark astrocytes, and found that the majority (~95%) of the GAL4+ glia are negative for GAT (Fig S1). In summary, *SPARC-GAL4*, *GRM56F03-GAL4*, and *GMR10E12-GAL4* mark ensheathing glia around the antennal lobe in the late pupal stage.

**Fig. 1.**
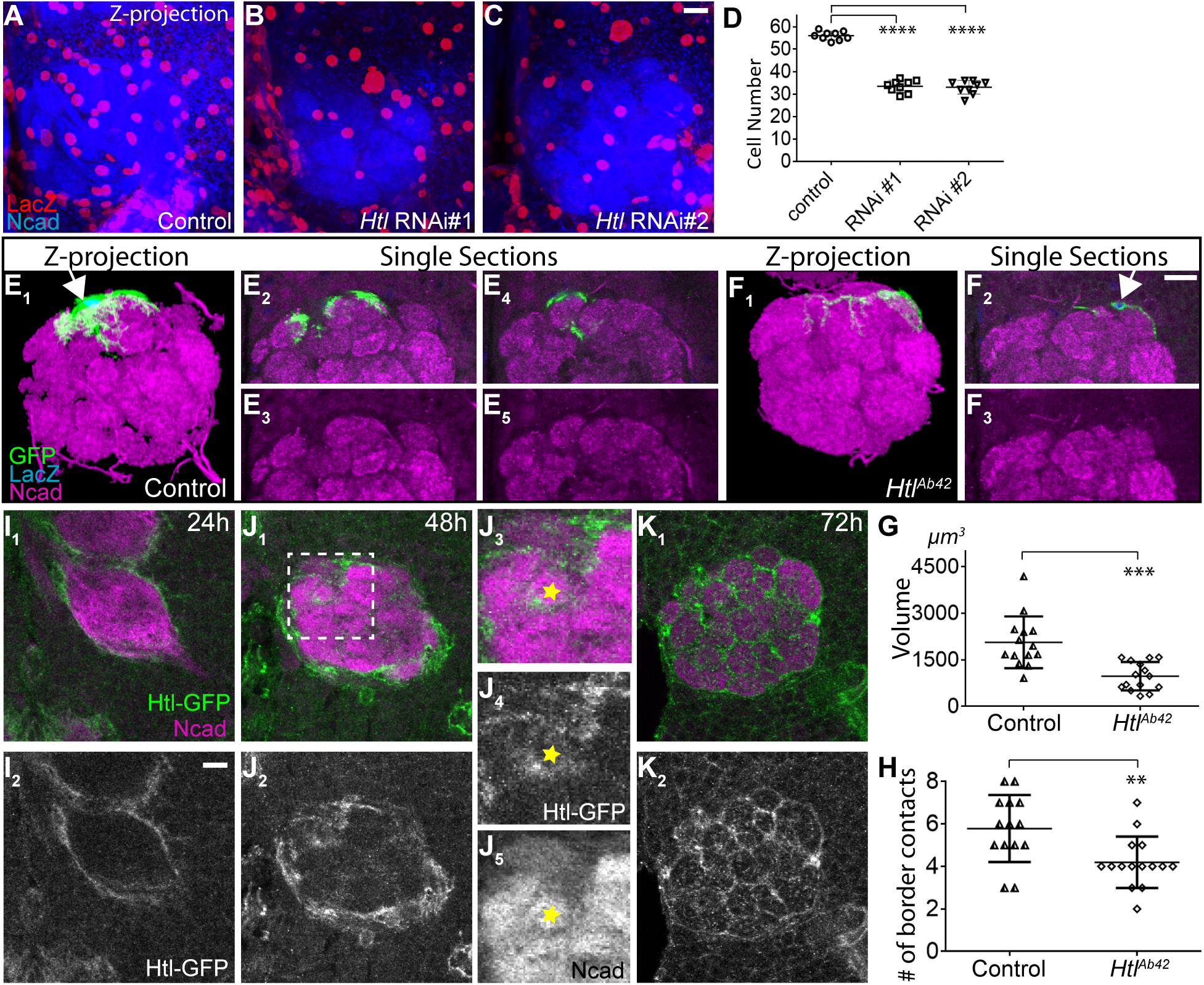
Ensheathing glia morphogenesis during antennal lobe development. *(A–D)* Confocal sections for antennal lobe at 24 h (A), 48 h (B), 72 h (C), and 96 h (D) APF. Ensheathing glia processes are labeled by *SPARC-GAL4* driven (>) *UAS-mCD8GFP*. Ensheathing glia nuclei are marked by *UAS-nuclear-LacZ*. Magenta shows neuropil staining by the N-cadherin (Ncad) antibody. Dashed rectangles are enlarged on the right column. D, dorsal; L, lateral. Scale bars are 10 μm. *(E)* Confocal sections of 96 h antennal lobe. Green shows glia nuclei stained by the Repo antibody, overlaid on the LacZ channel in *E_2_*. *(F)* Quantification of antennal lobe ensheathing glia cell number at different developmental stages, and the number of total glia around the antennal lobe at the end of pupae stage. Error bars represent standard deviation (SD). N=5 for each time point.

*SPARC-GAL4* also drove reporter expression at 24 h APF, which enabled us to investigate the development of antennal lobe ensheathing glia. At 24 h APF, prior to glomerular formation, ensheathing glia processes were restricted to the periphery of the antennal lobe (Fig 1A). From 24 h to 48 h APF, axon terminals of ORNs invade the antennal lobe and target specific subregions to connect with their synaptic partners such as dendrites from PNs (17). We found that around 48h APF when proto-glomeruli first emerged, ensheathing glia processes started to invade the antennal lobe from the lobe surface (Fig 1B). Even at this initial stage of infiltration, ensheathing glia processes displayed a preference to grow along the borders between adjacent proto-glomeruli, instead of growing into them (Fig 1B_3_). At 72 h APF, as glomerular structures became more clearly separable, ensheathing glia processes had encircled most glomeruli (Fig 1C). These processes remained on glomerular boundaries, with minimal extension into the glomeruli. Ensheathing glia wrapping of olfactory glomeruli was complete by the end of the pupal stage (Fig 1D).

We also observed an increase in the number of *SPARC-GAL4*+ ensheathing glia around the antennal lobe from 24 h to 72 h APF (Fig 1F). This increase is at least in part due to glial cell proliferation, since we could generate glia clones via the MARCM technique (27) with heat shock induced mitotic recombination at any point from 24 to 72 h APF (see Fig 3 and Fig S3 below).

### *Heartless* Knockdown in Ensheathing Glia Reduces Processes and Disrupts the Ensheathment Pattern

To identify the molecular signals that control the extension of ensheathing glia processes, we tested ~50 candidate adhesion molecules, receptor kinases, and receptor phosphatases by RNA interference (RNAi) and screened for potential defects in ensheathing glia development and antennal lobe organization. We found that expression of two independent RNAi targeting non-overlapping regions of *heartless* (*htl*), a fibroblast growth factor receptor (28-30), caused a 70% reduction of ensheathing glia processes within the antennal lobe (Fig 2A-D). This phenotype is consistent with a previous discovery that FGF signaling is required for astrocytes to infiltrate the larval ventral nerve cord of *Drosophila* (26). However, the antennal lobe provides a unique opportunity to observe the response of neuronal compartmentalization to glia morphogenesis defects.

**Fig. 2.**
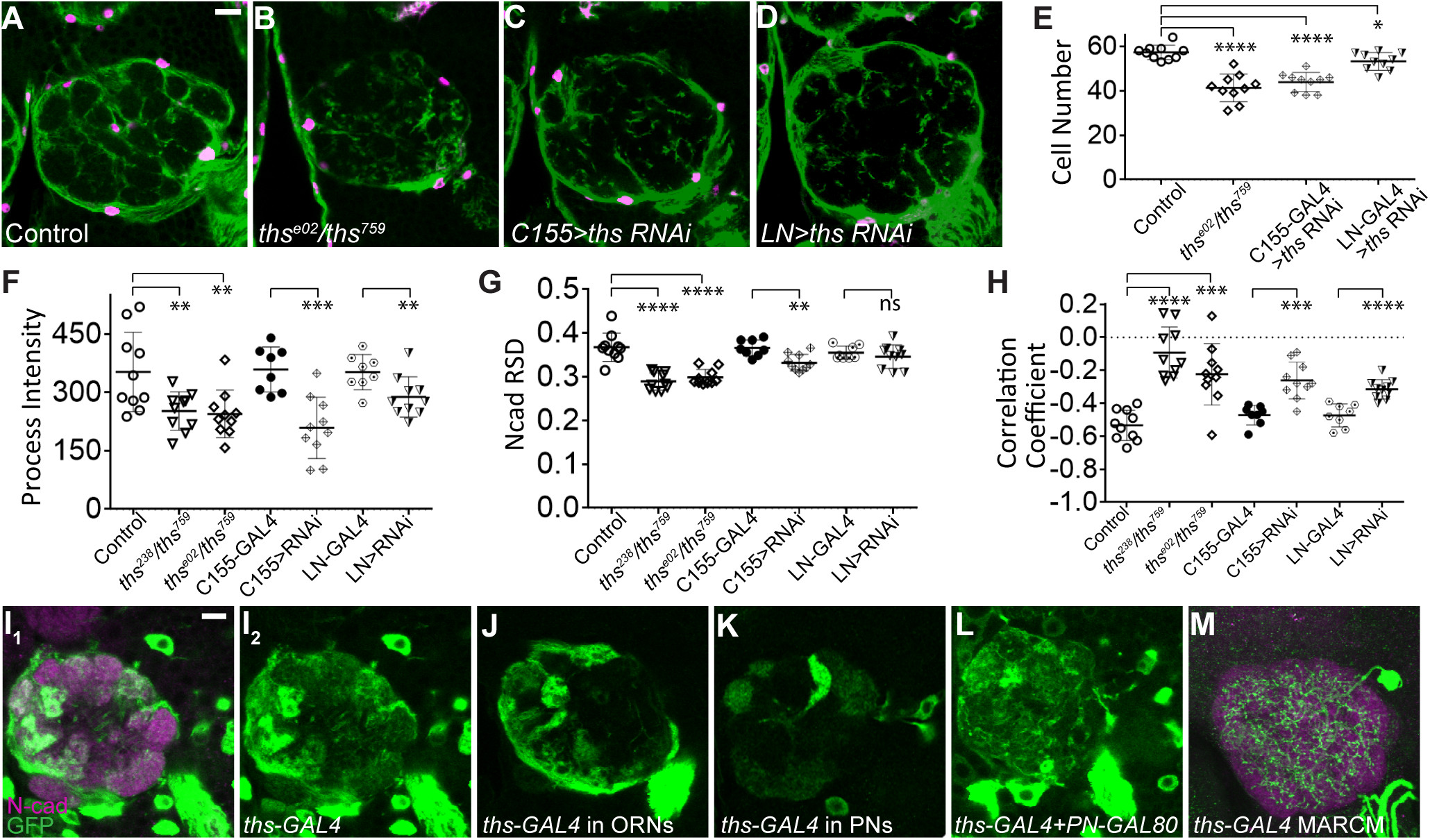
Heartless knockdown causes an altered ensheathing glia wrapping pattern and disrupts antennal lobe compartmentalization. *(A–C)* Confocal sections for the antennal lobe at 96 h APF of wild type (A) and two different *UAS-Htl RNAi* constructs (RNAi#1: VDRC6692, RNAi#2: VDRC27180) driven by *SPARC-GAL4 (B, C)*. Ensheathing glia processes are labeled by *SPARC-GAL4> UAS-mCD8GFP*. Neuropil compartments are stained with Ncad antibody. GFP and Ncad intensities along a line between the center of DM6 and DA1 glomeruli (indicated by dashed circles in *A_2_* and asterisks in *A_2_–C_2_*) are plotted in *A_3_–C_3_*, with the X-axis indicating the position of each point on the line from DM6 to DA1 (left to right); fluorescence intensity quantified in arbitrary unit. *(D)* Quantification of total GFP intensities normalized by antennal lobe size for wild-type and RNAi expressing flies. *(E)* Quantification of relative standard deviation of Ncad intensities for each antennal lobe of wild-type and RNAi flies. (F) Correlation coefficient of GFP and Ncad intensities as plotted in *A_3_–C*_3_. Scale bar is 10 μm. Error bars represent SD. **, p<0.01; ***, p<0.001, ****, p<0.0001.

We found that many glomeruli became less clearly separable (Fig 2A_2_, B_2_, C_2_, arrowheads) when *htl* was knocked down. To quantify this effect, we used relative standard deviation (RSD) of the N-cadherin neuropil staining intensity as an index for the degree of compartmentalization, as incomplete antennal lobe compartmentalization would cause obscure glomerular borders and hence smaller variance in the neuropil staining signal. We found a significant reduction of RSD in *htl* knockdown antennal lobes compared with wild type (Fig 2E). The remaining glomerular borders sometimes also lacked ensheathing glia processes that normally would divide the adjacent glomeruli (Fig 2A-C). We also observed ensheathing glia processes that extended within glomeruli, which is rarely observed in wild type antennal lobes (Fig 2B_1_, arrowhead). This is likely due to poor establishment of glomerular borders. We quantified the localization pattern of the ensheathing glia processes relative to the glomerular compartments by plotting the intensity of the GFP signal derived from glial membranes together with the intensity of Ncadherin signal marking the neuropil (Fig 2A_3_-C_3_), and calculated their correlation coefficient (Fig 2F). In control animals, N-cadherin and GFP signals exhibited strong anti-correlation (Fig 2F), since ensheathing glia processes are preferentially located on the borders of neuropil compartments instead of within the glomeruli. However, such anti-correlation was greatly diminished when *htl* was knocked down (Fig 2F), suggesting that loss of *htl* results in a disrupted ensheathment pattern.

### *Heartless* Promotes Ensheathing Glia Survival and Cell-autonomously Regulates Process Elaboration

Heartless is involved in cell differentiation, directional migration, and survival in a wide variety of tissues (26, 28, 31-34). Consistent with this notion, we observed that *htl* knockdown caused a 40% reduction of ensheathing glia number near the antennal lobe as assayed by the nuclear-LacZ marker (Fig 3A-D). This decrease is likely due to cell death, since the expression of an apoptosis suppresser, P35 (35), in ensheathing glia largely rescued the reduction of cell number (Fig S2); however, P35 expression did not rescue the reduction of ensheathing glia processes or the defect in glomerular compartmentalization (Fig S2), suggesting that FGF signaling may also regulate morphogenesis of antennal lobe ensheathing glia in addition to its role in supporting glia survival.

**Fig. 3.**
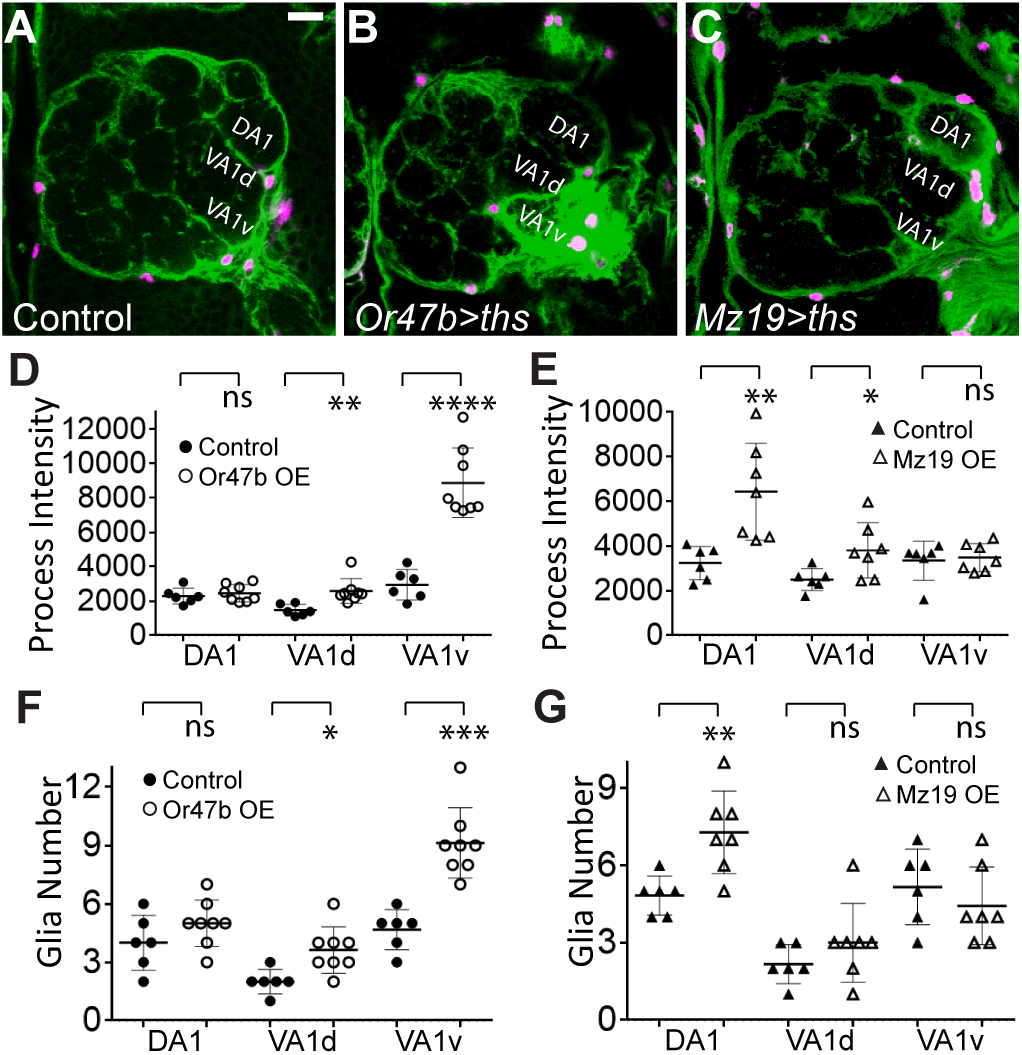
Heartless controls ensheathing glia number and cell-autonomously control their morphology. *(A–C)* Projections of confocal sections along the Z-axis. Red: ensheathing glia cell bodies marked by *SPARC-GAL4> UAS-nuclear-LacZ.* Blue: Ncad antibody stains for antennal lobe neuropil. *(D)* Quantification of ensheathing glia cell number in each antennal lobe. ****, p<0.0001. *(E, F)* Wild-type *(E)* and *htl^Ab42/Ab42^ (F)* ensheathing glia single cell MARCM clones in adult antennal lobe. Green shows ensheathing glia processes labeled by *GMR10E12-GAL4> UAS-mCD8GFP*. Blue shows ensheathing glia nuclei marked by *UAS-nuclear-LacZ*. Magenta shows Ncad antibody staining for antennal lobe neuropil. *(G)* Quantification of the volume of processes from each ensheathing glia labeled by MARCM. *(H)* Quantification of the number of borders each MARCM-labeled ensheathing glia contacts. Error bars represent SD. **, p<0.01; ***, p<0.001. (I–K) Confocal sections of antennal lobe at 24 h *(I)*, 48 h *(J)*, and 72 h *(K)* APF with Htl-GFP signal. Dashed rectangle in *J_1_* is enlarged and shown as *J_3_–J_5_*. The yellow stars mark the center of a proto-glomerulus around which ensheathing glia is wrapping. Scale bars are 10 μm.

To dissociate the role of *htl* in controlling glia survival and process extension, we used the MARCM technique (27) to generate single-cell clones of ensheathing glia homozygous for the *htl^Ab42(null)^* (30) allele in an otherwise heterozygous background. In this experiment, *GMR10E12-GAL4* was used to label the homozygous mutant cells. We used *UAS-mCD8GFP* to visualize glial processes (Fig 3E-F), and *UAS-nuclear-LacZ* (Fig 3E1, F2, arrows) to visualize cell bodies of ensheathing glia in these MARCM clones. Compared to wild type, *htl^Ab42^* single-cell clones exhibited a 50% reduction of the volume of glial processes (Fig 3G) and a 50% reduction of total fluorescence intensity (Fig S3). These reductions were accompanied by a decrease in the number of glomeruli each ensheathing glia could access, as quantified by the number of glomerular borders each glia extended to (Fig 3H). Thus, this mosaic experiment demonstrated that *htl* is cell-autonomously required for ensheathing glia process extension.

To determine the subcellular localization of the Htl protein, we used a fosmid transgenic line that produces Htl-GFP from an insertion that contains the extended genomic region covering *htl* (36) to analyze GFP signal within and around the antennal lobe during pupal development. At 24 h APF, no Htl-GFP signal was detected inside the antennal lobe (Fig 3I), consistent with the location of ensheathing glia processes (Fig 1A). At 48 h APF, we started to detect Htl-GFP signal in between proto-glomeruli (Fig 3J). By 72 h APF, Htl-GFP signal was detected at most glomerular borders, though a lower level of signal was also detected within glomeruli (Fig 3K), which may originate from cell types other than ensheathing glia, such as astrocytes and potentially neurons. The signal on glomerular borders coincided well with the ensheathing glia wrapping pattern over this developmental course (Fig. 1). These data support the hypothesis that Htl mediates FGF signaling to regulate ensheathing glia process invasion of the antennal lobe and wrapping of individual glomeruli.

### Thisbe, an FGF ligand, Is Required for Ensheathing Glia Wrapping of Glomeruli

We next assessed the identity and cellular source of FGF ligands that regulate ensheathing glia development. Htl responds to two FGF ligands, Thisbe (Ths) and Pyramus (Pyr) (37, 38). To test whether *ths* is required for antennal lobe ensheathing glia to form the wrapping pattern, we analyzed ensheathing glia and antennal lobe morphology in animals that carried different *ths* mutant alleles (39). The phenotypes we observed with loss of *htl* were all recapitulated in trans-heterozygous combinations of *ths* alleles (*ths*^e02026^/*ths*^759^ and *Df(2R)ED2238*/*ths*^759^). There was an overall reduction of ensheathing glia processes and cell number in the antennal lobe of trans-heterozygous flies (Fig 4A-B, E-F). Similar to loss of *htl* from ensheathing glia, the antennal lobe of *ths* mutants also showed defective compartmentalization as measured by reduced RSD values of N-cadherin staining (Fig 4G) and a markedly reduced anti-correlation between the intensity of ensheathing glia processes and neuropil signal (Fig 4H). These findings indicate that *ths* is required for ensheathing glia to wrap around glomeruli in the antennal lobe, and that failure to form a correct wrapping pattern could disrupt compartmentalization of antennal lobe neuropil.

**Fig. 4.**
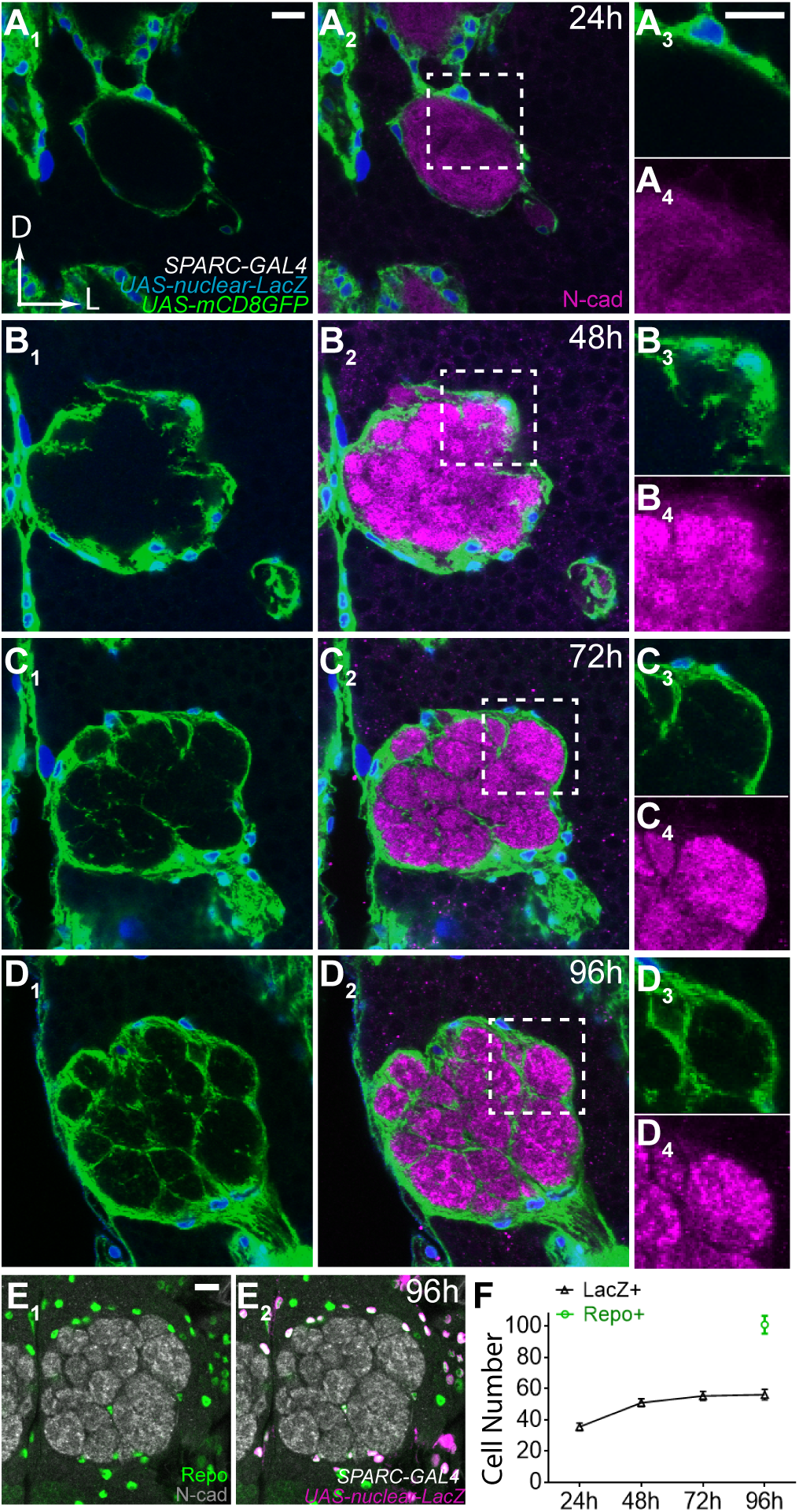
Thisbe is expressed in olfactory neurons and required for ensheathing glia to wrap glomeruli. *(A–D)* Ensheathing glia wrapping pattern in wild type *(A)*, ths mutant *(B; ths^e02^* denotes *ths^e02026^* allele), panneural C155-GAL4 (C) and LN *orb^0449-GAL4^* (D) driven RNAi against ths. Green shows ensheathing glia processes labeled by *SPARC-QF> QUAS-mtdT*. Magenta shows ensheathing glia nuclei marked with *QUAS-nuclear-LacZ*. *(E–H)* Quantification of ensheathing glia number *(E)*, process intensity (F), relative standard deviation of Ncad staining *(G)*, and the correlation coefficient for the intensities of ensheathing glia process and neuropil staining *(H)*. Error bars represent SD. *, p<0.05; **, p<0.01; ***, p<0.001; ****, p<0.0001; ns, p>0.05. *(I) ths* expression pattern revealed by ths-GAL4> UAS-mCD8GFP (green). Magenta in I1 shows Ncad counterstaining. *(J) ey-FLP* intersects with ths-GAL4 together with *UAS-FRT-stop-FRT-mCD8GFP* to show ths expression pattern in ORNs. *(K) GH146-FLP* intersection shows ths expression pattern in PNs. *(L) ths-GAL4+* LN cell bodies and LN and ORN processes, after *GH146-GAL80* suppressing *ths-GAL4* in most PNs. *(M)* MARCM labels a single LN that is positive for *ths-GAL4*. Green: GFP. Magenta: Ncad. All images are confocal sections of adult antennal lobe except panel (M), which is a projection of Z stacks. Scale bars are 10 μm.

### Thisbe Is Produced by ORNs, PNs, and Antennal Lobe Local Interneurons (LNs)

To identify which cell types express Thisbe, we took advantage of a MiMIC insertion (23) located between two coding exons of the *ths* gene. We converted the MiMIC cassette to an artificial exon that contains the coding sequence for *2A-GAL4* (40). Thus, GAL4 can be produced along with the endogenous N-terminus of Ths (encoded by the first two exons), and can drive reporter gene expression in the *ths* pattern. We then used this *Ths-GAL4* to express *UAS-mCD8GFP* to visualize the cell bodies and projections of the Ths-producing cells. In the antennal lobe, neuronal processes from PNs, ORNs, and LNs are all labeled, suggesting that all three major neuronal types produce Ths (Fig 4I).

To determine the contributions of each of these cell types, we used an intersectional strategy, in which *Ths-GAL4* was combined with Flp recombinases that are specifically expressed in ORNs (*ey-FLP*) (41) or PNs (*GH146-FLP*) (42). With a FLP-out reporter, *UAS-FRT-stop-FRT-mCD8GFP* (42), we found a number of ORN and PN classes were *Ths-GAL4*+ based on glomerular labeling (Fig 4J-K). At least 45 cell bodies over all sections of the antennal lobe were *Ths-GAL4+* without intersection (Fig 4I); however, only 19 of them were *GH146-Flp*+ PNs (which constitutes the majority of PNs innervating ~40 glomeruli), consistent with the small number of glomeruli (~7) labeled by intersection of *GH146-Flp* and *Ths-GAL4* (Fig 4K). This difference is likely due to the contribution from LNs, whose cell bodies are also located around the antennal lobe (43, 44), whereas ORN cell bodies are located in the peripheral sensory organs.

We used two approaches to validate that LNs produce Thisbe. First, we suppressed the expression from most PNs by combining *GH146-GAL80* (42) together with *Ths-GAL4*. After PN-derived signal was largely eliminated, around 27 cell bodies around the antennal lobe remained. The projection pattern of these neurons covered the entire antennal lobe (Fig 4L), which is characteristic of the majority of LNs (43). Second, we used *Ths-GAL4* to label single cells by the MARCM technique. We were able to identify MARCM clones for ORNs and PNs (data not shown), as well as LNs (Fig 4M). In summary, Ths is produced by a subset of ORNs, PNs, and LNs.

### LN-derived Thisbe Is Necessary for Ensheathing Glia Wrapping

To test in which cell type(s) Thisbe functions to regulate ensheathing glia morphogenesis and antennal lobe compartmentalization, we used RNAi to knock down *ths* in ORNs (*Pebbled-GAL4*), PNs (*GH146-GAL4*), LNs, and all neurons (*C155-GAL4*), respectively, while labeling the ensheathing glia by *SPARC-QF*, which we converted from *SPARC-GAL4* (23). Pan-neuronal knockdown of *ths* recapitulated the phenotypes observed in *ths* mutants (Fig 4C, E-H). Knocking down *ths* specifically in LNs by *orb^0449-GAL4^* (Fig. S4), a GAL4 line identified from the InSite screen (45), resulted in a mild but significant defect in ensheathing glia wrapping and antennal lobe glomerulus integrity (Fig 4D, E-H). Knocking down *ths* in PNs did not cause significant defects (Fig S5). We have also used MARCM combined with a cell lethal strategy (41) to create a near pan-ORN mutant background for *ths*, and did not find defects in ensheathing glia wrapping (Fig S5). The milder phenotypes in LN knockdown compared to pan-neuronal knockdown could be explained by: 1) pan-neuronal GAL4 may have stronger and/or earlier expression than the LNGAL4, and therefore causes more effective knockdown; 2) LN-GAL4 does not include all LNs; 3) Ths from ORNs and PNs synergize with Ths from LNs. Regardless, our data indicate that LN is an essential cellular source of Ths in the antennal lobe for directing ensheathing glia wrapping.

### Thisbe Can Instruct Glomerular Wrapping with a High Spatial Specificity

We have shown that FGF signaling is necessary for ensheathing glia to extend processes into the antennal lobe and demarcate individual glomeruli. To test whether FGF signaling can instruct ensheathing glia to wrap around selected neuropil compartments, we expressed *ths* in only one class of ORNs (VA1v) using a GAL4 under the control of the promoter of the odorant receptor specifically expressed in this class (*Or47b-GAL4*). We observed that overexpression resulted in hyper-wrapped VA1v glomerulus by ensheathing glia (Fig 5B, D). This effect was highly localized, since in addition to VA1v, only the adjacent VA1d glomerulus was slightly hyper-wrapped (Fig 5D), likely due to intensified ensheathing glia processes on its border shared with VA1v. Hyper-wrapping did not extend to the DA1 glomerulus (Fig 5D), which is one glomerulus away from the VA1v glomerulus. There was also an excess of glia cells around the hyper-wrapped VA1v glomerulus (Fig 5F).

**Fig. 5.**
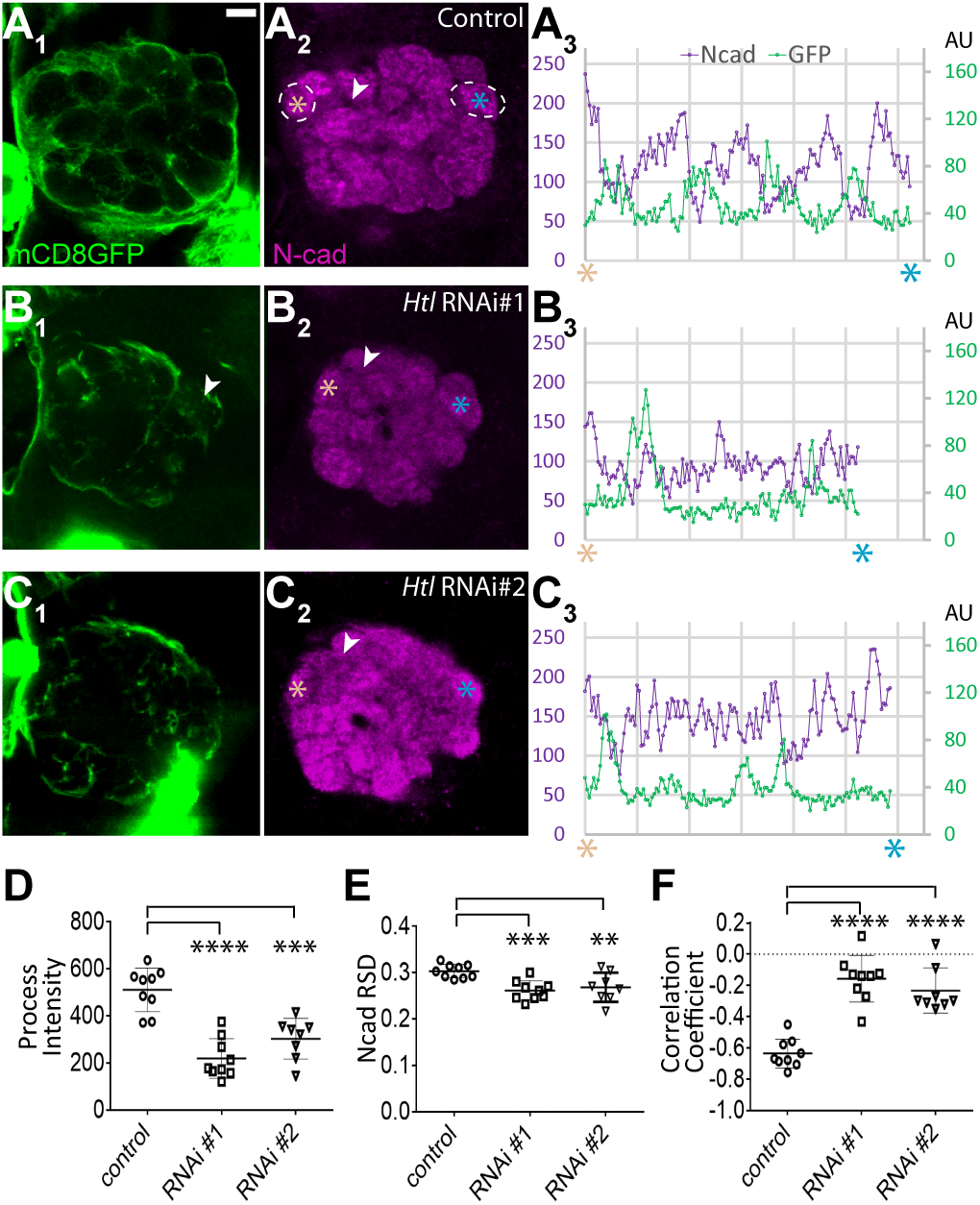
Changes in ensheathing glia processes and number in response to local overexpression of ths. *(A–C)* Confocal sections of adult antennal lobe of wild type *(A)*, overexpression of *UAS-ths* by *Or47b-GAL4 (B)* or by *Mz19-GAL4 (C)*. Green: *SPARC-QF>QUAS-mtdT* labels ensheathing glia processes. Magenta: Ensheathing glia nuclei marked with *QUAS-nuclear-LacZ*. Scale bar is 10 μm. *(D, E)* Quantification of signal intensities of ensheathing glia processes around each glomerulus normalized by the perimeter of the corresponding glomerulus. *(F, G)* Quantification of ensheathing glia numbers within 5-μm distance to the surface of each glomerulus. Error bars represent SD. *, p<0.05; **, p<0.01; ***, p<0.001; ****, p<0.0001; ns, p>0.05.

Similarly, overexpressing Ths in *Mz19-GAL4*+ PNs, which send dendrites to DA1 and VA1d, also caused local hyper-wrapping of these glomeruli, as well as a local increase of ensheathing glia cells (Fig 5C, E, G). We have consistently noticed that *Mz19-GAL4* activity is stronger in DA1 PNs than in VA1d PNs (46). Accordingly, glial hyper-wrapping was more pronounced around DA1 than around VA1d. These results suggest that Thisbe acts locally as a spatial cue to instruct ensheathing glia to infiltrate the antennal lobe.

### FGF Signaling Ensures Accurate Neuronal Targeting

Lastly, we examined whether glia wrapping defects could disrupt neuronal projections to the glomerular compartments. We used membrane proteins under the control of specific odorant receptor promoters (*Or88a-mtdT* and *Or47b-rCD2*) to label two ORN classes that project their axons to two adjacent glomeruli, VA1v and VA1d (Fig 6A). When *ths* was knocked down with a pan-neuronal driver, *C155-GAL4*, VA1v and VA1d ORN axons still reached their target area in the ventrolateral antennal lobe. However, they failed to establish exclusive territories for their axonal arborization, and instead had partially overlapped axonal terminals (Fig 6B). Likewise, in *ths* mutant animals, VA1v and VA1d axon terminals partially intermingled, rather than confining to their respective glomerular compartments (Fig 6C; quantified in Fig 6D). Thus, FGF signaling between neurons and ensheathing glia is essential for antennal lobe compartmentalization and targeting accuracy for ORN axons.

**Fig. 6.**
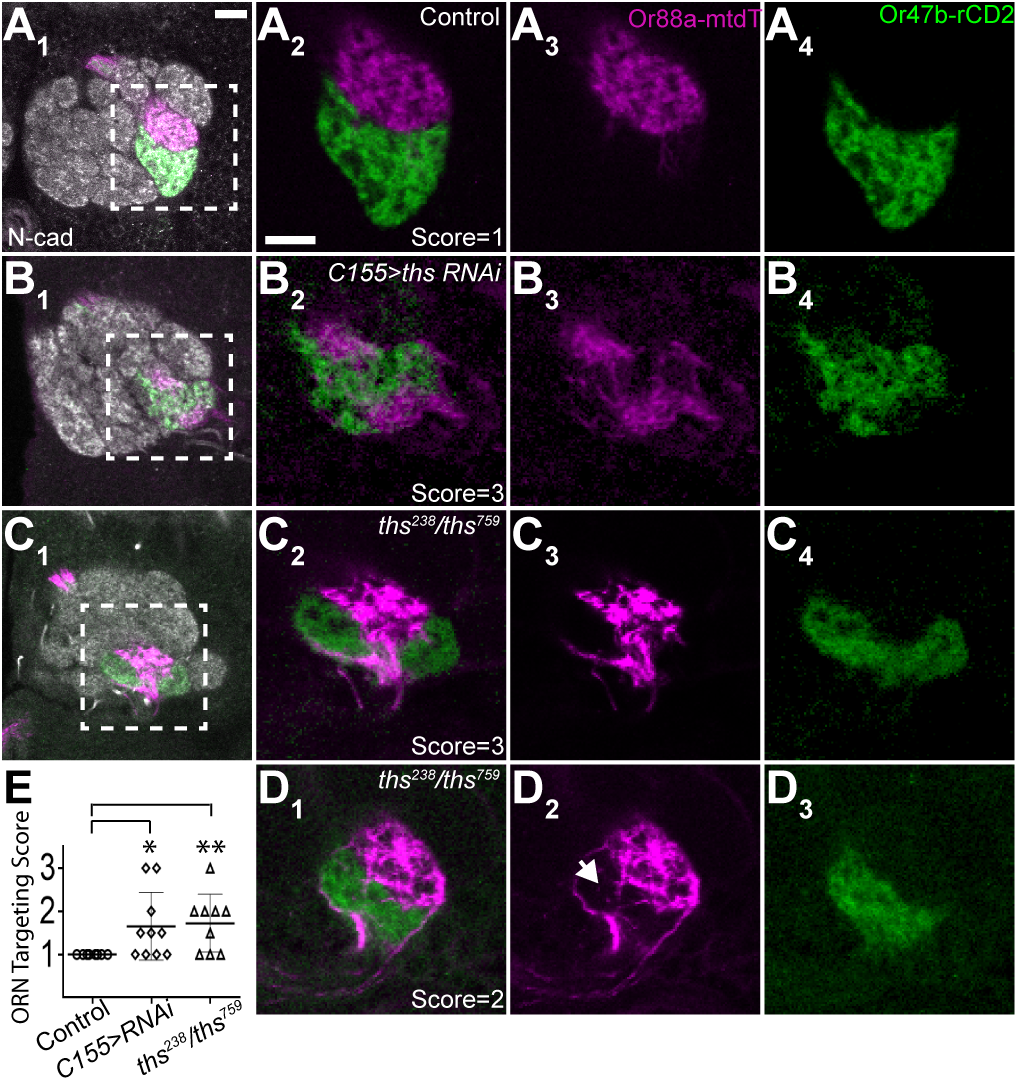
Loss of ths results in olfactory receptor neuron targeting defects. *(A–D)* Confocal sections of the adult antennal lobe, with *Or88a-mtdT* and *Or47b-rCD2* labeling ORN axons that target VA1d (magenta) and VA1v (green) glomerulus, respectively. Images within the dashed rectangles in the first column are enlarged in columns 2–4. *(A)* Wild type. *(B) C155-GAL4* driven *ths RNAi*. *(C-D)* ths mutant flies showed severe intermingling of VA1d and VA1v ORN terminals (C) and a mild spillover of VA1d ORN axons (D, arrow). *ths^238^* denotes the *Df(2R)ED2238* allele. Scale bars are 10 μm. *(E)* Quantification of the accuracy of ORN axon targeting. 1, normal; 2, mild spillover; 3, severe intermingling. Each symbol represents 1 fly with two antennal lobes scored separately and averaged. Error bars represent SD. *, p<0.05; **, p<0.01.

## Discussion

The use of discrete neuropil compartments for organizing and signaling information is widespread in invertebrate and vertebrate nervous systems. In both the fly antennal lobe and vertebrate olfactory bulb, axons from different ORN classes are segregated into distinct glomeruli (47). The rodent barrel cortex also uses discrete compartments, the barrels, to represent individual whiskers (48). In this study, we show that FGF signaling between neurons and glia mediate neural compartment formation in the *Drosophila* antennal lobe.

Members of the FGF family have diverse functions in a variety of tissues in both vertebrates and invertebrates (49, 50). Vertebrate FGFs regulate not only neural proliferation, differentiation, axon guidance, and synaptogenesis, but also gliogenesis, glial migration, and morphogenesis (51-55). Many of these roles are conserved in invertebrates. For example, Thisbe and Pyramus induce glial wrapping of axonal tracts (32, 33) much like the role of other FGF members played in regulating myelin sheaths in mammals (55). Thisbe and Pyramus also control *Drosophila* astrocyte migration and morphogenesis (26). Likewise, FGF signaling also promotes morphogenesis of mammalian astrocytes (56). Therefore, studying the signaling pathways in *Drosophila* will extend our understanding of the principles of neural development.

In ensheathing glia, whose developmental time course and mechanisms have not been well documented prior to this study, we observed a glial response to FGF signaling reminiscent of the paradigm shown before (26, 32, 57); however, the exquisite compartmental structure of the *Drosophila* antennal lobe and genetic access allowed us to further scrutinize changes of neuropil structure and projection patterns that occurred alongside morphological phenotypes in ensheathing glia. We demonstrated a necessity of Ths in local interneurons, although it is possible that ORNs and PNs also contribute. We also tested the function of the other ligand, Pyramus, in antennal lobe development. We did not detect any change in ensheathing glia morphology with *pyramus* RNAi, and double RNAi against *thisbe* and *pyramus* did not enhance the phenotype compared to *thisbe* knockdown alone (data not shown).

FGF signaling in glomerular wrapping appears to be highly local. In our overexpression experiments, the hyper-wrapping effect was restricted to the glomerulus where the ligand is excessively produced, and did not spread to nearby non-adjacent glomeruli. These experiments suggest that Thisbe communicates locally to instruct glial ensheathment of the glomeruli, rather than diffusing across several microns to affect nearby glomeruli. Since heparan sulfate proteoglycans are known to act as FGF co-receptors by modulating the activity and spatial distribution of the ligands (50, 58, 59), we speculate that Thisbe in the antennal lobe may be subject to such regulation to limit its diffusion and long-range effect.

Our data showed that deficient ensheathment of antennal lobe glomeruli is accompanied by imprecise ORN axon targeting. However, we cannot determine whether these targeting defects reflect initial axon targeting errors, or a failure to stabilize or maintain the discrete targeting pattern. Previous models for the establishment of antennal lobe wiring specificity suggested that the glomerular map is discernable by the time glia processes start to infiltrate the antennal lobe (17). Due to a lack of class-specific ORN markers for early developmental stages, it is still unclear the relative timing between when neighboring ORN classes refine their axonal targeting to discrete compartments and when ensheathing glia barriers are set up. Regardless, our discovery that FGF signaling functions in the formation of discrete neuronal compartments in the antennal lobe highlights an essential role for glia in the precise assembly of neural circuits.

## Methods

### Immunostaining

Tissue dissection and immunostaining were performed according to previously described methods (60). Primary antibodies used in this study include rat anti-DNcad (DN-Ex #8; 1:40; DSHB), chicken anti-GFP (1:1000; Aves Labs), rabbit anti-DsRed (1:500; Clontech), mouse anti-rCD2 (OX-34; 1:200; AbD Serotec), rat anti-HA (1μg/ml, Roche), mouse nc82 (1:35; Developmental Studies Hybridoma Bank, [DSHB]), mouse anti-Repo (1:50, DSHB), rabbit anti-β-Galactosidase (MP Biomedicals, 1:125), rabbit anti-GAT (1:3000; a gift from Marc Freeman), and mouse anti-β-Galactosidase (1:1000; Promega). Secondary antibodies were raised in goat or donkey against rabbit, mouse, rat, and chicken antisera (Jackson Immunoresearch), conjugated to Alexa 405, FITC, 568, or 647. Confocal images were collected with a Zeiss LSM 780 and processed with Zen software, ImageJ, and Imaris.

### Mosaic Analysis

The hsFlp MARCM analyses were performed as previously described (27, 61) with slight modifications. *GMR10E12-GAL4* was used for labeling ensheathing glia in adult stage. Flies were kept at 18°C and were heat shocked for 30min at 37°C between 0-24h APF.

### Data Analysis

Confocal sections of antennal lobes (Fig. 2 and 4) were analyzed by ImageJ software for measuring integrated intensity value, area size, mean, and standard deviation for the region of interest (ROI) manually selected based on N-cadherin counterstaining. Process intensity was calculated using the integrated intensity value of the antennal lobe section crossing DA1 and DM6 glomeruli normalized by the area size of the antennal lobe of that section. The Plot Profile function in ImageJ was used to measure signal intensities for ensheathing glia processes and Ncad along the selected lines. Glia cell number was manually counted based on LacZ staining within 10-μm distance to the surface of the antennal lobe determined by Ncad counterstaining. To quantify the MARCM clones (Fig. 3), confocal sections of marked ensheathing glia were processed by Imaris software through manually thresholding the images by a set of consistent parameters, followed by automatic measurement of the volume and total intensity of glia processes. In the overexpression experiment (Fig. 5), ROI was manually selected by including the inner region and the boundaries of the glomerulus of interest based on Ncad counterstaining. Process intensity was defined as the integrated intensity value divided by the perimeter of the ROI selection. Glia number was manually counted based on LacZ signal within 5-μm distance to the surface of each glomerulus. ORN targeting defects (Fig. 6) were scored by an experimenter blind to the genotype. Graphs were generated using the GraphPad Prism software. Mean ± SD were shown on the graphs. Statistical significance was calculated with Graphpad Prism using two-tailed Student’s t test (Fig. 2-5) or Mann-Whitney test (nonparametric data in Fig. 6).

## Acknowledgments

We thank M. Freeman, C. Klämbt, H. Bellen, T. Clandinin, and Vienna and Bloomington Stock Centers for reagents, X. Wang, J. Lui, A. Shuster, J. Ren, H. Li, T. Li, and L. DeNardo for discussion and helpful comments on the manuscript. This work was supported by an NIH grant (R01 DC005982) to L.L., who is an HHMI investigator.

